# Sodium chloride is an ionic checkpoint for human Th2 cell responses and shapes the atopic skin microenvironment

**DOI:** 10.1101/390393

**Authors:** Julia Matthias, Julia Maul, Rebecca Noster, Hanna Meinl, Ying-Yin Chao, Heiko Gerstenberg, Florian Jeschke, Gilles Gasparoni, Anna Welle, Walter Jörn, Karl Nordström, Klaus Eberhardt, Dennis Renisch, Sainitin Donakonda, Percy Knolle, Dominik Soll, Stephan Grabbe, Natalie Garzorz-Stark, Kilian Eyerich, Tilo Biedermann, Dirk Baumjohann, Christina E. Zielinski

## Abstract

Sodium is an ionic checkpoint for the induction and amplification of human Th2 cell responses and shapes the atopic skin microenvironment, where it could serve as a novel therapeutic target for Th2 mediated diseases.

**Abstract:** There has been a strong increase in the incidence of allergic diseases over the last 50 years. Environmental factors most likely account for this phenomenon. However, the nature of these factors and the mode of action by which they induce the type 2 immune deviation, which is characteristic of atopic diseases, remains unclear. It has previously been reported that dietary sodium chloride promotes the polarization of Th17 cells with implications for autoimmune diseases such as multiple sclerosis. Here, we demonstrate that sodium chloride also potently promotes Th2 cell responses on multiple regulatory levels. Sodium chloride enhanced IL-4 and IL-13 production while suppressing IFN-*γ* production in effector T cells. It diverted alternative T cell fates into the Th2 cell phenotype and also induced *de novo* Th2 cell polarization from naïve T cell precursors. Mechanistically, it exerted its effects via the osmosensitive transcription factor NFAT-5 and the kinase SGK-1, which regulated Th2 signature cytokines and master transcription factors in hyperosmolar salt conditions. The skin of patients suffering from atopic dermatitis contained highly elevated amounts of sodium compared to non-lesional atopic and healthy skin. This demonstrates that sodium chloride represents a so far overlooked cutaneous microenvironmental factor in atopic dermatitis that can induce Th2 cell responses, the orchestrators of allergic diseases. Together, our data propose ionic signaling through sodium chloride as a novel checkpoint and potential therapeutic target for type 2 immunity and its associated allergic diseases.

## Introduction

Naïve T helper cells can differentiate into functionally diverse subsets of effector T cells that are tailored to distinct antimicrobial responses. They integrate polarizing cues such as TCR signal strength, costimulation and cytokine signaling in order to differentiate into distinct T cell subsets (*1*). Still, much remains unknown with respect to how signals are recognized and integrated to determine cell fate and function. Microenvironmental factors in the tissues are thought to provide additional cues that drive T helper cell differentiation. Recently, ionic checkpoints such as sodium chloride (NaCl) and potassium have been demonstrated to modulate T cell responses. In particular, NaCl has been shown to boost Th17 cell differentiation from naïve T cell precursors with implications for the pathogenesis of multiple sclerosis under high-salt diet conditions (*2–4*). Moreover, elevated levels of potassium following tumor cell necrosis have been shown to provide a tumor permissive tissue microenvironment by paralyzing cytotoxic T cell functions (*5*). Therefore, the ionic composition of the microenvironment represents a novel, yet poorly investigated determinant of T cell polarization and T cell effector functionalities.

Epidemiological studies provide robust support for a fast increase in the incidence of allergic and autoimmune diseases over the last 50 years. This is thought to be due to environmental non-genetic changes of yet unclear origin (*6*). More than one hundred years ago, a renowned pediatric reference book claimed the improvement of atopic dermatitis upon dietary salt restriction based on clinical observations but without any mechanistic evidence (*7*). Considering the steady increase in the dietary consumption of processed food and thus of NaCl over the last 50 years, we set out to investigate a potential link between NaCl mediated ionic signaling and enhanced Th2 cell responses, which are the culprits in the pathogenesis of allergic diseases.

Our data demonstrate the existence of an ionic determinant of Th2 immunity, which challenges the current cytokine-focused concept of T cell polarization. NaCl induced and enhanced Th2 cell responses via the transcriptional target NFAT-5 and the kinase SGK-1, independently of IL-4. In line with the Th2 promoting effects of NaCl, we could also demonstrate enhanced NaCl deposition in the skin of patients suffering from atopic dermatitis, a Th2 driven skin disease. Together, our data propose ionic signaling through NaCl as a novel checkpoint and thus potential therapeutic target for type 2 immunity in allergic diseases.

## Results

### Sodium chloride enhances Th2 cell effector functions

Recent reports have demonstrated a strong impact of sodium chloride (NaCl) on boosting Th17 cell differentiation from naïve T cell precursors in polarizing cytokine conditions (*2, 3*). However, NaCl is particularly enriched in peripheral barrier tissues such as the skin, which naïve T cells do not enter during their regular recirculation route (*8*). Therefore, we sought to determine whether NaCl could also regulate T cell responses on the memory and effector T cell level, which would support a role of ionic signals for *in situ* immunomodulation.

We purified CD4^+^ CD45RA^−^ memory T cells and analyzed IL-17 expression on the single-cell level following expansion with CD3 and CD28 antibodies in the absence (low NaCl) or presence (high NaCl) of additional 50 mM NaCl, which reflects physiological NaCl concentrations in peripheral blood and skin, respectively (*9*). Further increase of NaCl beyond additional 50 mM decreased cell viability and resulted in cell death (**Fig. S1**). NaCl strongly enhanced IL-17 expression in memory T helper cells in the absence of exogenous polarizing cytokines (**Fig. 1A**). It also increased IL-22, a Th17 associated cytokine involved in the repair of barrier tissues as well as other Th17 cell associated signature molecules such as the master transcription factor ROR-*γ*t, the chemokine receptor CCR6 as well as the anti-microbial Th17 cytokine IL-26 (**Fig. S2**) (*10–12*).

**Figure 1.**
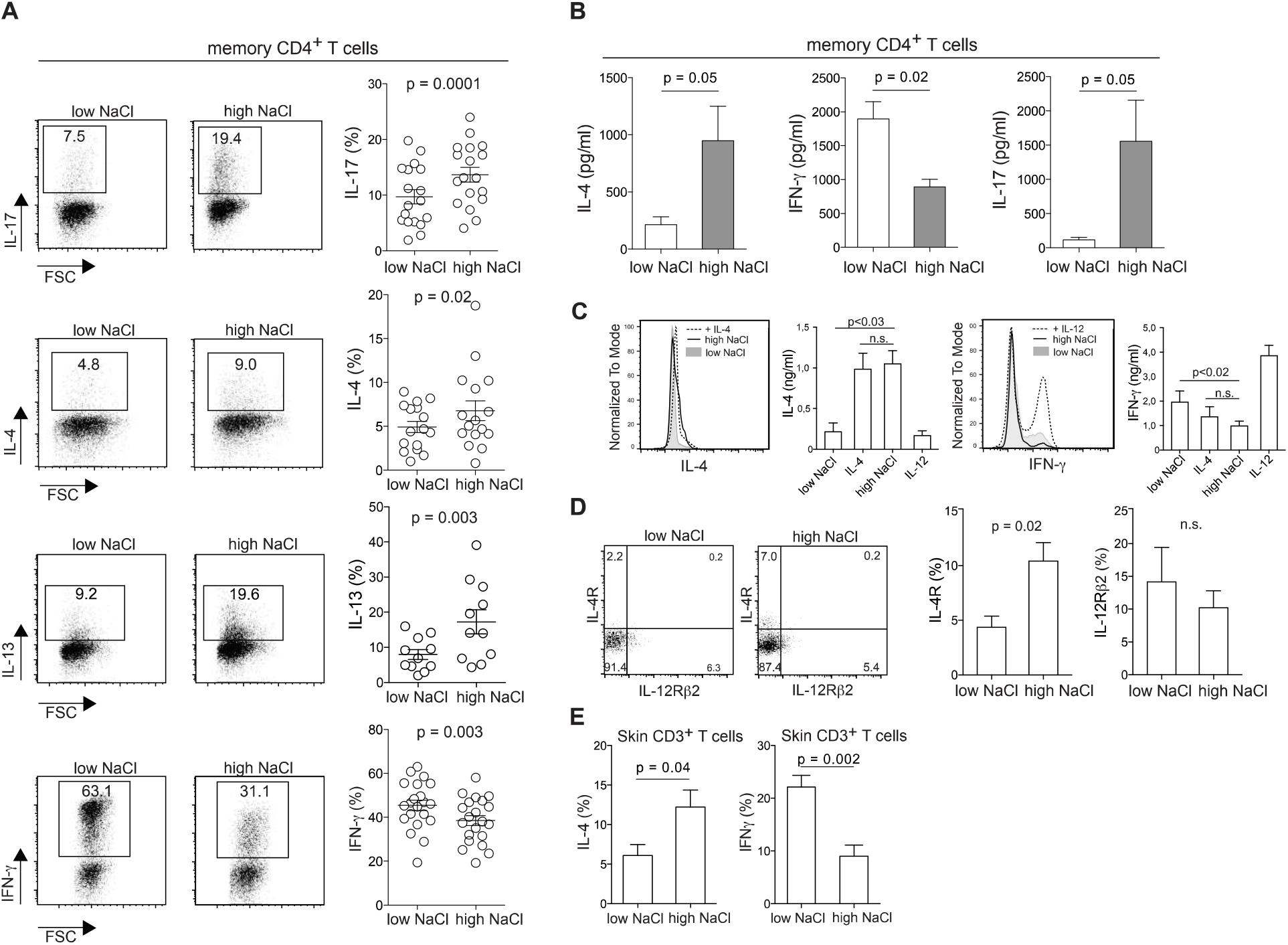
Sodium chloride enhances Th2 and suppresses Th1 cell responses in memory T cells. (A-D) Human memory T helper cells were sorted from fresh PBMC as CD4^+^CD14^−^ CD45RA^−^ T cells by flow cytometry and stimulated for a total culture period of 5 days in the presence (high) or absence (low) of additional 50 mM NaCl (compared to baseline cell culture medium including supplements) with CD3 and CD28 mAb for 48h. (**A**) Intracellular staining and flow cytometry (FACS) on day 5 following PMA and ionomycin restimulation. Shown is a FACS staining of an individual experiment (left panel) and cumulative data with each circle representing one donor. (**B**) ELISA of cell culture supernatants analyzed on day 5 (n = 3). (**C**) Flow cytometry and ELISA as in A-B to compare the relative effect of NaCl to recombinant IL-4 and IL-12 (FACS panels are representative of three experiments; ELISA, n = 3). (**D**) Flow cytometry for the cytokine surface receptors IL-4R and IL-12β2. Shown is one representative experiment and a summary graph of cumulative data (n = 5). (**E**) CD3^+^ T cells were sorted by flow cytometry after enzymatic digestion (collagenase IV 0,8mg/ml, 16 hours) of fresh human skin obtained after abdominoplasties. T cells were characterized by flow cytometry according to **Fig. S4** and stimulated and analyzed as in (**A**) (n = 3).

Since memory T cells display a complex cytokine expression pattern, we also studied the response of other T helper cell lineage defining cytokines to NaCl. Surprisingly, we observed upregulation of the Th2 signature cytokine IL-4 by both intracellular cytokine staining and ELISA, even in the presence of IL-4 neutralizing antibodies (**Fig. 1A**, **Fig. 1B**, **Fig. S3**, **Fig. S4**). The Th2 signature was corroborated by IL-13 upregulation (**Fig. 1A**). IFN-*γ* expression was downregulated in accordance with the reciprocal regulation of Th2 and Th1 pathways. This Th2 bias was further supported by the upregulation of IL-4, IL-13 and IL-5 and downregulation of IFN-*γ* in T helper cells clones upon restimulation in high NaCl conditions (**Fig. S5**). Of note, NaCl acted as an equally efficient signal for Th2 cytokine upregulation and IFN-*γ* suppression as recombinant IL-4, which has so far been considered to be the principal Th2 polarizing factor (**Fig. 1C**) (*13*). NaCl also upregulated IL-4R expression on memory T helper cells, which supports a feedback amplification of the Th2-axis (**Fig. 1D**). IL-12Rβ2 expression remained unaffected. The Th2-promoting effect was restricted to ionic osmolytes containing sodium and was strongest for NaCl among all tested tonicity signals. A Th2 skewing could not be induced by non-ionic osmolytes such as urea or mannitol, which were tested at various concentrations (**Fig. S6A, B**).

We also extended our investigations on the impact of NaCl signals to the CD8 T cell compartment. IL-17 was significantly upregulated in CD8 memory T cells under elevated NaCl conditions, although to a much lower extent than in CD4 memory T cells. IL-4 production, however, was not increased in CD8 T cells in contrast to CD4 T cells (**Fig. S7**).

We next tested the response of T cells isolated from healthy human skin to restimulation with NaCl. Peripheral tissues contain heterogeneous memory T cell populations, which differ from those in the blood (*14*). Their functional plasticity and response to immunomodulatory factors remain poorly defined. Skin resident T cells are particularly prone to exposure to ionic signals exerted by fluctuating levels of NaCl, since the skin is known to act as a reservoir for excess dietary NaCl (*15–17*). To characterize the identity of skin T cells, we phenotyped them according to their differential expression of CD69 and CD103, as both markers correlate with tissue residency (*18*). This confirmed the heterogeneous composition of skin memory T cells, which is distinct from memory T cell subsets in the blood (**Fig. S8**). We found robust IL-4 upregulation with concomitant IFN-*γ* downregulation in skin resident T cells upon stimulation with NaCl (**Fig. 1E**). In addition, we also observed upregulation of the skin homing chemokine receptor CCR8 by NaCl, which correlates with higher IL-4 and lower IFN-*γ* expression levels, as well as with skin T cell residency (**Fig. S9**) (*19, 20*). Therefore, NaCl also exerts Th2 promoting effects on cutaneous memory T cell subsets, which reside at sites of elevated NaCl levels *in vivo.*

Overall, these findings demonstrate that ionic signals exerted by NaCl can directly alter human memory and effector T cell functions in the absence of exogenous cytokine signals. NaCl has very potent Th2 promoting functions that match the Th2 polarizing potential of IL-4.

### Sodium chloride induces transcriptional activation of the Th2 program

To further corroborate that NaCl can promote the Th2 identity, we analyzed the expression of master transcription factors in response to T cell stimulation with NaCl (**Fig. 2**). GATA-3, the master transcription factor of Th2 cells, was highly upregulated in memory T cells upon T cell receptor stimulation in the presence of additional NaCl, whereas T-bet, the master transcription factor of Th1 cells, was downregulated (**Fig. 2A**). We also generated T cell clones to avoid the selective outgrowth of undefined T cell subpopulations and subjected each one individually to restimulation in the presence or absence of additional NaCl. GATA-3 was highly upregulated in memory T cell clones upon NaCl stimulation, while T-bet was downregulated, again supporting the Th2-promoting role of NaCl (**Fig. 2B**). In accordance with higher IL-17 expression levels (**Fig.1A**, **Fig. S5B**), ROR-*γ*t expression was also enhanced upon NaCl stimulation (**Fig.2B**).

**Figure 2.**
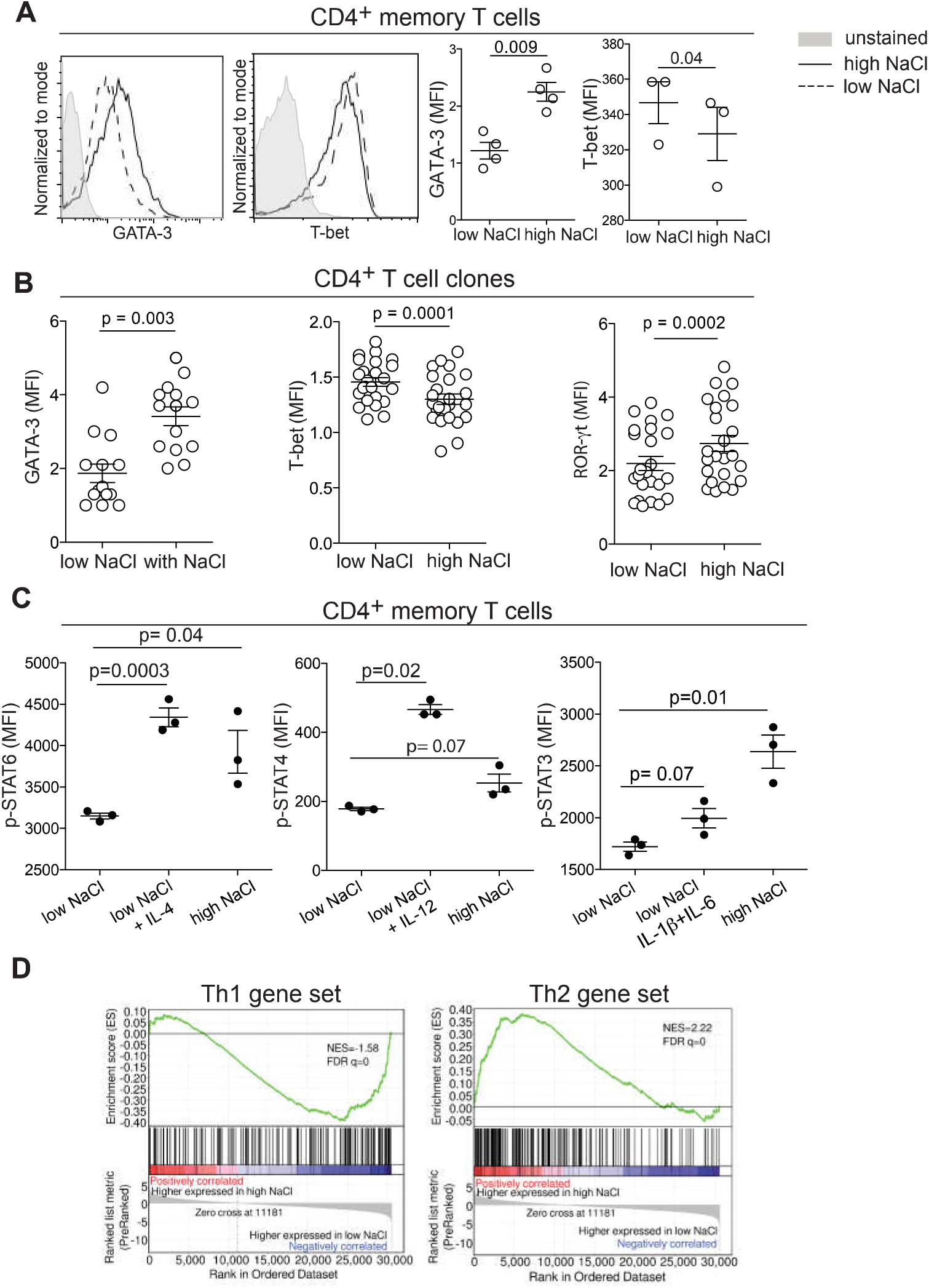
Sodium chloride induces the transcriptional activation of Th2 and suppression of Th1 programs. (**A**) Human memory T cells (CD4^+^CD45RA^−^) were stimulated with CD3 and CD28 mAb for 48h of a 5-day culture period in the presence (high) or absence (low) of additional 50 mM NaCl before intracellular staining for transcription factors on day 5. Shown are representative flow cytometry analyses (left panel) and cumulative data (right panel) with each circle indicating one donor and experiment. (**B**) T cell clones were generated from CD4^+^ CD45RA^−^ memory T cells during a 14-day culture period with irradiated allogeneic feeder cells and phytohemagglutinin and were restimulated and analyzed as in (**A**). T cell clones were randomly selected from growing cultures for restimulation experiments in low or high NaCl conditions. Each circle represents an individual T cell clone. (**C**) CD4^+^CD45RA^−^ memory T cells were stimulated as in (A-B) in low or high NaCl conditions or in the presence of recombinant polarizing cytokines in low NaCl conditions for 5 days before restimulation for 30 min with p-STAT6 (IL-4), p-STAT4 (IL-12) and p-STAT3 (IL-6) inducing cytokines, respectively. Phosphorylation of STAT molecules was assessed after intracellular staining and flow cytometry. (**D**) The GSEA enrichment plots of the transcriptome of CD4^+^CD45RA^−^ memory T cells treated in low or high NaCl conditions compared to Th1 (left panel) and Th2 (right panel) gene sets. The green curve of the plots shows the running enrichment score. NES, normalized enrichment score, FDR q values represent the statistical significance of enrichment.

Signal transducer and activator of transcription proteins (STATs) have crucial roles in transmitting cytokine signals and in specifying T helper cell polarization (*21*). For that reason, we investigated the effect of NaCl on lineage-specifying STAT molecules (**Fig. 2C**). Memory T helper cells were stimulated polyclonally with anti-CD3 and anti-CD28 in the presence or absence of additional NaCl for 5 days or with polarizing cytokines such as IL-4, IL-12, or IL-6 and IL-1β. On day 5, T cells were washed and restimulated with IL-4 or IL-12 for 30 minutes to induce STAT6 or STAT4 phosphorylation, respectively. T cells that were stimulated in the presence of NaCl displayed stronger STAT6 phosphorylation upon IL-4 exposure than control T cells cultured in the absence of exogenous NaCl. STAT4 phosphorylation, however, remained unaffected by NaCl stimulation. STAT3 was strongly upregulated by NaCl in memory T helper cells (**Fig. 2C**), in line with its IL-17 promoting properties (*22, 23*). STAT3 is also known to bind to Th2-associated gene loci and to cooperate with STAT6 for Th2 induction, further supporting the NaCl-induced Th2 bias (*24, 25*). Together, these results argue that NaCl promotes the Th2 cell signature also on the level of the transcriptional STAT gate keepers.

In order to provide a more global view on the T helper cell polarization bias induced by NaCl, we performed next-generation mRNA sequencing to analyze the gene expression changes of memory T helper cells stimulated in the presence of low versus high NaCl concentrations. Gene set enrichment analysis (GSEA) of the differentially expressed genes using Th1 and Th2 associated gene sets corroborated the strong Th2 bias also on a transcriptome-wide level (**Fig. 2D**).

### Sodium chloride skews distinct T helper cell subsets to acquire Th2 properties

We next wanted to determine if distinct T helper cell fates displayed plasticity to adopt the Th2 phenotype upon exposure to NaCl. We therefore isolated fully differentiated human Th1, Th2 and Th17 cell subsets *ex vivo* from peripheral blood according to their differential expression of chemokine receptor surface markers, which, according to our previous work, reliably identifies the respective T helper cell lineages (*11, 23*). After 5 days of stimulation with CD3 and CD28 mAb, the T cell subsets maintained their characteristic signature cytokine profiles as expected. In the presence of additional NaCl, Th2 cells further upregulated IL-4, while their low baseline IFN-*γ* production was further downregulated, promoting their Th2 identity. Interestingly, Th2 cells also significantly upregulated IL-17, although absolute IL-17 expression levels remained low. Th1 cells did not alter their low IL-4 or IL-17 production upon restimulation with NaCl, but significantly down-regulated IFN-*γ*. Of note, Th17 cells also increased IL-4 production, while downregulating IFN-*γ* and increasing IL-17 (**Fig. 3A**). To corroborate plasticity as the underlying process to acquire Th2 properties by non-Th2 cells, we generated T cell clones and subjected individual clones to restimulation in the presence of additional NaCl. This approach confirmed that homogenous clonal T cell populations could acquire and increase IL-4 production, while at the same time expressing decreased levels of IFN-*γ* and increased levels of IL-17 on the clonal level (**Fig. 3B**). Importantly, we also observed that IL-4-negative T cell clones could acquire the ability to produce IL-4 in response to restimulation with NaCl (**Fig. 3B, C**). This *de novo* acquisition of cytokine expression also applied to individual IL-17 negative T cell clones, which upregulated IL-17 expression upon restimulation with NaCl (**Fig. 3C**).

**Figure 3.**
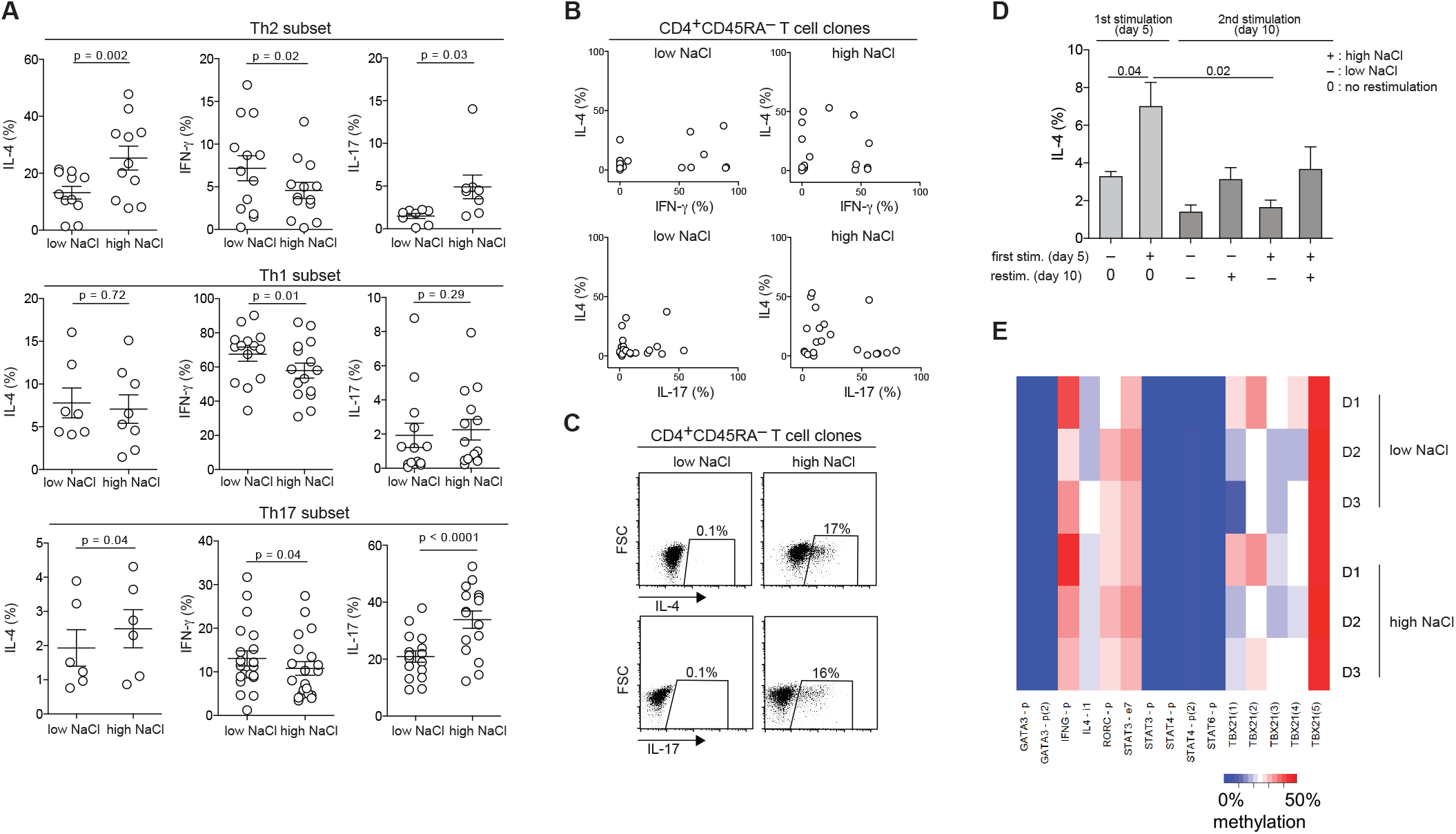
Sodium chloride reprograms distinct T helper cell subsets to acquire Th2 properties. (**A**) Polarized human T helper cell subsets were sorted *ex vivo* according to the differential expression of chemokine receptor surface markers as Th1 (CXCR3^+^CCR4^−^CCR6^−^), Th2 (CXCR3^−^CCR4^+^CCR6^−^) and Th17 cells (CXCR3^+^CCR4^−^CCR6^+^) and restimulated in the presence (high) or absence (low) of additional 50 mM NaCl with CD3 and CD28 mAb for 48h of a 5-day culture period. Cytokine expression was determined by intracellular cytokine staining and flow cytometry following PMA and ionomycin restimulation for 5 h. (**B**) Random T cell clones that were generated from CD4^+^CD45RA^−^ memory T cells were stimulated as in (**A**) and their cytokine coexpression pattern tracked in the presence of low versus high NaCl concentrations. Every circle represents a unique T cell clone. (**C**) Individual T cell clones that were generated from CD4^+^CD45RA^−^ memory T cells were selected based on being negative for IL-4 or IL-17 expression. They were restimulated in the presence of low versus high NaCl concentrations for 5 days before intracellular staining following PMA and ionomycin restimulation. IL-4 and IL-17 expression levels following NaCl treatment were highly variable. Shown are two individual clones from one experiment and blood donor (n = 3). (**D**) Human CD4^+^CD45RA^−^ memory T cells were sorted by flow cytometry and stimulated for 48h with CD3 and CD28 mAb for a total culture period of 5 days in low or high NaCl conditions before intracellular cytokine staining and flow cytometric analysis following PMA and ionomycin stimulation on day 5. The same T cell cultures were then split and restimulated for another 5 days in low and high NaCl conditions before intracellular cytokine staining and flow cytometric analysis following PMA and ionomycin stimulation on day 10. n = 4. (**E**) No DNA methylation differences in Th2 cells from three different donors cultured with (+) or without additional (-) NaCl. Heatmap shows average CpG methylation as measured by deep sequencing of bisulfite PCR amplicons for indicated regions. (p = promoter, I, intron, e = exon, numbers in brackets indicate amplicons for different subregions of the respective genes, *: paired t-test p-value = 0.045).

Since memory T cells are long-lived and can be reactivated in changing microenvironmental contexts by their cognate antigens, we investigated the stability of NaCl induced functional changes in human T helper cells. To this end, we stimulated memory T helper cells in high NaCl conditions for 5 days before transferring them into low NaCl conditions for another 5 days of anti-CD3 and anti-CD28 restimulation, and vice versa. The data demonstrate that the upregulation of IL-4 is dependent on the presence of high NaCl concentrations since these increased IL-4 expression levels could not be maintained upon restimulation in low NaCl conditions. Repetitive restimulation compromised overall IL-4 production. This was partially counteracted if restimulation occurred in high NaCl conditions (**Fig. 3D**).

It has previously been shown that epigenetic mechanisms regulate the stability and plasticity of T cell programs (*26*). We therefore determined whether ionic signaling by NaCl could epigenetically imprint the Th2 program in human T helper cells. To this end, we analyzed DNA methylation signatures of selected regulatory regions at several candidate loci, including STAT genes (*STAT3, STAT4 and STAT6), RORC, GATA3, Tbx21, IL4,* and *IFNG* in response to NaCl using targeted ultra-deep sequencing of bisulfite converted DNA. Our data suggest that the increase in Th2 effector functions is not exerted by methylation changes at the analyzed regions (**Fig. 3E**, Table S1). Together, NaCl mediated ionic signaling most likely preserved the flexibility of T cell effector functions, in line with the functional re-adaptation to low NaCl conditions shown in the restimulation experiments (**Fig. 3D**).

### NaCl promotes Th2 cell differentiation from naïve T cell precursors in the absence of polarizing cytokines

Given the Th2-promoting effects of NaCl on memory and effector T cell subpopulations, we next sought to determine whether it could exert direct polarizing effects on the naïve T cell compartment. Although NaCl has previously been demonstrated to promote human Th17 cell differentiation, this only occurred in the presence of exogenous polarizing cytokines (*2*). We therefore isolated human naïve T cells to high purity and stimulated them in the presence or absence of additional NaCl. Interestingly, we observed a significant increase of IL-4 and IL-13 production upon stimulation with NaCl in the absence of polarizing cytokines (**Fig. 4A**). This also occurred in the presence of IL-4 neutralizing antibodies (**Fig. S3**). As expected, IFN-*γ* could be induced by polyclonal stimulation but was suppressed in the presence of exogenous NaCl, in line with the reciprocal Th1-Th2 regulation pattern. Also, IL-17 producing cells could be slightly, although significantly, induced by NaCl, even in the absence of exogenous polarizing cytokines, while polyclonal stimulation alone had no IL-17 promoting effect. These findings were supported by quantifying cytokine release by ELISA (**Fig. 4B**). Similar to memory cells, naïve cells selectively upregulated GATA-3 expression, but not T-bet and ROR-*γ*t, upon NaCl stimulation (**Fig. 4C**). Under Th17 polarizing conditions (IL-1β + IL-6 + TGF-β + IL-23 + IL-21), we observed an abrogation of NaCl induced IL-4 upregulation in human naïve T cells (**Fig. S10**). This cytokine microenvironment masked the direct Th2 promoting effect of NaCl that we have revealed here and is therefore in line with previous work that has missed to detect the induction of Th2 cell responses upon NaCl stimulation in Th17 polarizing conditions (*2*).

**Figure 4.**
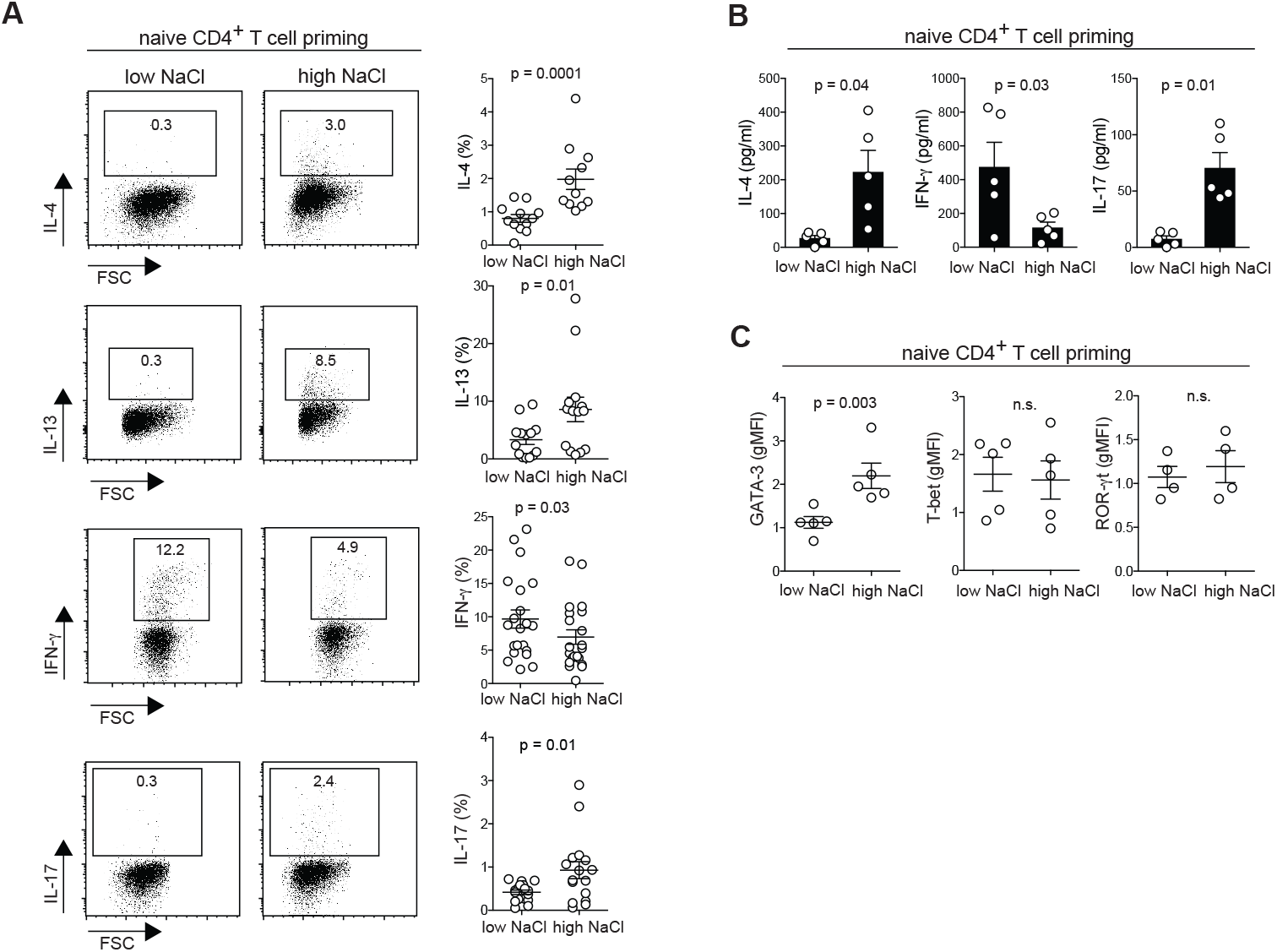
Sodium chloride promotes Th2 cell differentiation from naïve T cell precursors in the absence of polarizing cytokines. (A, C) Human naïve CD4^+^CD45RA^+^CCR7^+^ T cells were isolated by flow cytometry and stimulated for 48h with CD3 and CD28 mAb in the presence of low or high NaCl concentrations for a culture period of 5 days. Cytokine and transcription factor expression was determined by flow cytometry following PMA and ionomycin restimulation for 5h. (**A**) left panel, cytokine expression in one individual blood donor; right panel, cumulative data of all donors. (**B**) Supernatants from the cultures in (**A**), were tested by ELISA and normalized to cell numbers by counting beads. Each circle represents a separate blood donor.

To further corroborate a role for hypersalinity in the enhancement of type 2 immunity, we tested its effects on murine T cell polarization. Naïve T cells from C57BL/6 mice were stimulated with CD3 and CD28 mAb in the presence of distinct polarizing cytokines and cytokine blocking antibodies that are known to reliably polarize Th1, Th2, Th17 and Treg responses *in vitro.* Naïve T cell priming in the presence of additional 40 mM NaCl strongly enhanced Th2 cell priming with recombinant IL-4 with respect to IL-4 and GATA-3 expression. The presence of additional NaCl also induced slight IL-4 and GATA-3 upregulation in several non-Th2 cell conditions despite the presence of IL-4-antagonistic cytokines such as IL-12 and TGF-β (**Fig. S11**). Similar results were obtained with CD4^+^ T cells derived from BALB/c mice (not shown). Increased IL-4 concentrations were also detected in the culture supernatant of Th2 cells that were treated with additional NaCl (**Fig. S11D**).

Together, these findings demonstrate that NaCl acts as a Th2 cell polarizing signal during the priming of naïve T cells as assessed by multiple readouts in both humans and mice. Moreover, this occurred in the absence of exogenous Th2 polarizing cytokines such as IL-4 as well as independent of autocrine IL-4 signaling and, in mice, even despite the presence of Th2-counteracting cytokines.

### SGK-1 and NFAT-5 mechanistically link sodium chloride signaling with the Th2 program

Having established that NaCl has significant effects on the induction and enhancement of of the Th2 cell program, we sought to investigate the underlying molecular mechanisms. We tested whether the nuclear factor of activated T cells 5 (NFAT-5), which is known to be an osmosensitive transcription factor, is induced in human memory T helper cells upon polyclonal activation in the presence of NaCl (*27*). NFAT-5 was significantly upregulated as assessed on its mRNA transcript level (**Fig. 5A**). Serum-and glucocorticoid-regulated kinase 1 (SGK-1) is also known to be regulated by tonicity signals and to be a downstream target of NFAT-5 (*28*). Accordingly, shRNA-mediated silencing of *NFAT5* downregulated *SGK1* transcript levels in low and high NaCl conditions, whereas silencing of *SGK1* had not impact on *NFAT5* expression levels (**Fig. S12**). Similar to NFAT-5, SGK-1 was more abundant in CD3 and CD28 stimulated memory T helper cells in the presence of additional NaCl (**Fig. 5A**). Interestingly, we found that these osmosensitive molecules directly control the Th2 program in hyperosmolar conditions since short hairpin RNA (shRNA) induced silencing of NFAT-5 as well as SGK-1 abrogated NaCl-induced IL-4 upregulation as well as IFN-*γ* suppression (**Fig. 5B**). This was further corroborated by GATA-3 and T-bet expression levels, which were also downregulated by NFAT-5 or SGK-1 silencing to levels reflecting base-line ionic signaling (**Fig. 5C**). shRNAs directed against NFAT-5 and SGK-1 did not regulate IL-4 or IFN-*γ* production or GATA3 or TBX21 expression in low NaCl conditions (**Fig. 5B**, **Fig. S13**). Interestingly, this reveals that cytokine regulation can adopt distinct transcriptional pathways depending on the context of the tissue microenvironment, i.e. hyperosmolarity. Together, these data demonstrate a role of osmosensitive transcription factors for the regulation of the Th2 program in hyperosmotic tissue microenvironments.

**Figure 5.**
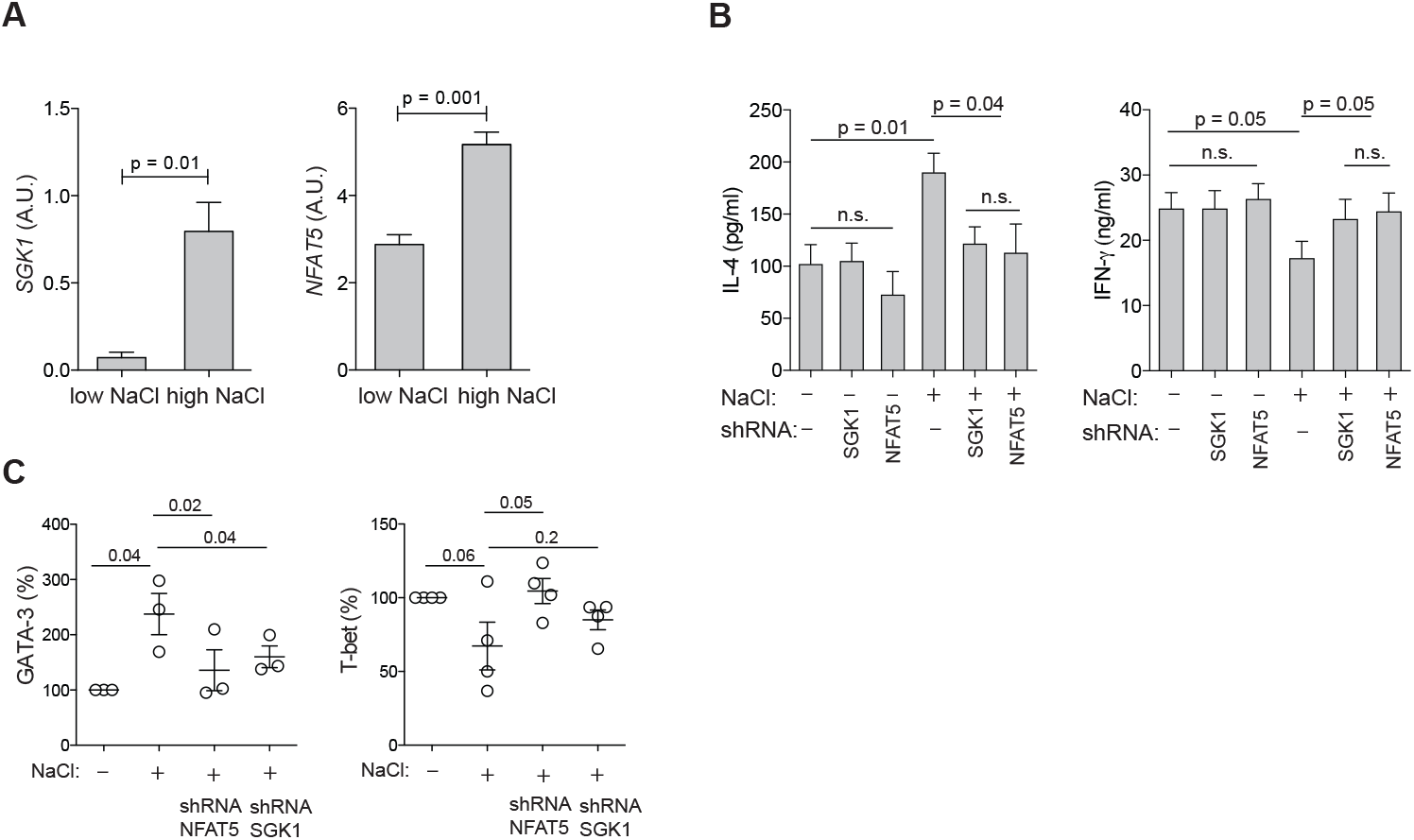
SGK-1 and NFAT-5 mechanistically link salt signaling with the Th2 program. (**A**) Human memory CD4^+^CD45RA^−^ T cells were stimulated with CD3 and CD28 mAb for 48h and cultured for 5 days in the presence (high) or absence (low) of additional 50 mM NaCl. SGK1 and NFAT5 expression were determined by αRT-PCR. (**B**) Cytokine production in supernatants was determined by ELISA after stimulation in low or high salt conditions for 5 days and shRNA-mediated silencing of SGK1 or NFAT5. (**C**) Intracellular staining and flow cytometry of master transcription factors after stimulation in low or high salt conditions for 5 days and shRNA-mediated silencing of NFAT5 and SGK1.

### Atopic dermatitis skin lesions display strongly increased sodium concentrations

Finally, we aimed to identify whether NaCl is associated with the pathogenesis of allergic diseases in humans. Despite intense and long-term research efforts, it still remains elusive how the Th2 bias is initiated in allergies (*29*). To this end, we investigated NaCl concentrations in the skin of patients suffering from atopic dermatitis (syn. atopic eczema), a Th2-driven chronic inflammatory skin disease with a high prevalence and strongly increasing incidence in industrialized nations (*30*). Neutron activation analysis (NAA) is very well suited for the detection and quantification of sodium in an organic matrix, since the mass elements in organic tissue like O, H, C, or N are hardly activated by thermal neutron irradiation, whereas sodium exhibits a high activation cross section and thus absorption probability for thermal neutrons and its activation product Na-24 emits two clearly measurable gamma lines for its identification (*31*). We therefore took advantage of neutron irradiation in a research neutron source to activate human tissue and to record gamma-spectra for element identification and determination of concentrations. The high penetrating ability of neutrons provides an effective means of probing elemental content of materials without prior sample preparation and manipulation and with the highest possible precision.

Lesional skin of patients suffering from moderate to severe atopic dermatitis (mean SCORAD = 63) displayed strongly increased concentrations of sodium compared to matched non-lesional patient skin (30-fold) (**Fig. 6A**, **Table S2**). We did not observe any difference in the sodium content between non-lesional skin of atopic dermatitis patients and skin from healthy donors. No sodium enrichment was detected in the inflamed skin of patients suffering from moderate to severe psoriasis (mean PASI = 25). This excludes that NaCl accumulation is a general phenomenon of the inflamed skin and supports its role for the pathogenesis of atopic dermatitis (**Fig. 6B**).

**Figure 6.**
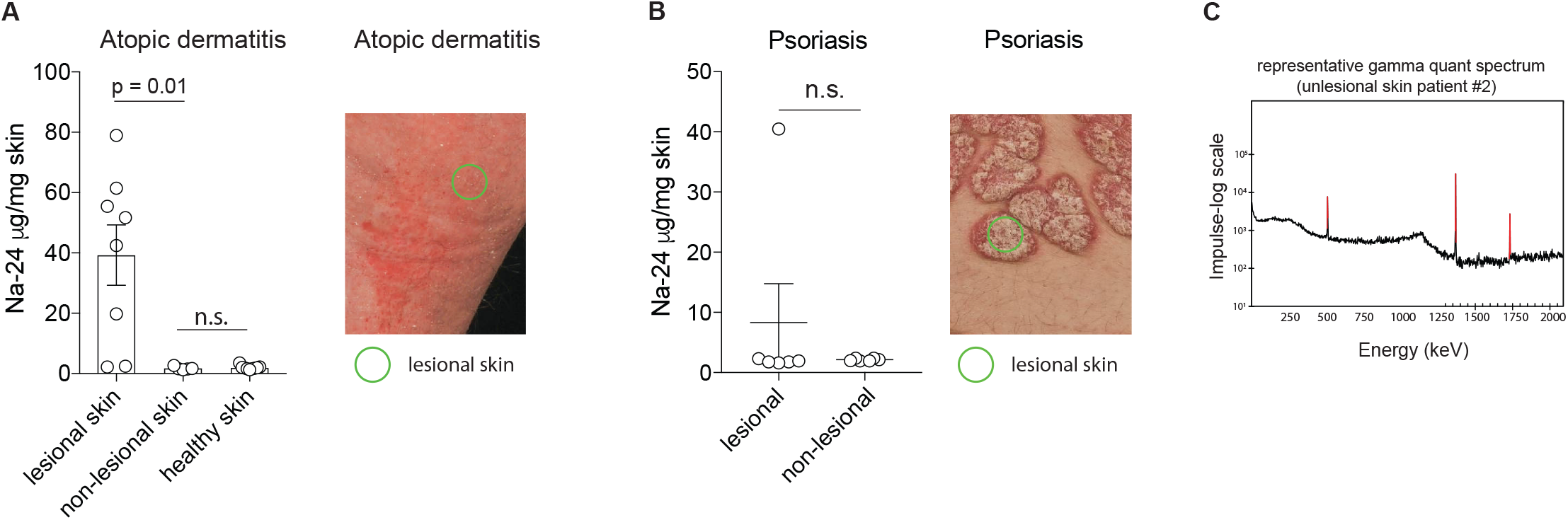
The skin of atopic dermatitis patients has a strongly increased sodium content. (**A**) Neutron activation analysis (NAA) of 4 mm skin punch biopsies was performed. The punch biopsies were taken from lesional and non-lesional skin of atopic dermatitis patients and of healthy controls (patients undergoing plastic surgery). Patient information is provided in Table 2. The concentration of sodium is shown (mean ± SEM). The right panel illustrates the clinical presentation of a representative patient with atopic dermatitis. (**B**) NAA of 4mm punch biopsies from lesional and non-lesional skin of psoriasis patients. The right panel illustrates the clinical presentation of a representative patient with psoriasis. (**C**) Shown is a representative spectrum of gamma quanta emission. The spectrum shows the accumulated counting rate of gamma quanta as a function of energy. The abscissa covers an energy range between 0 and 2 MeV. The prominent peaks at 1368 keV and 1732 keV are related to the decay of Na-24 and have been used to calculate the Na-concentration in the sample. The peak at 511 keV has to be attributed to the electron-positron annihilation radiation and cannot be used for the identification of the emitter. The spectrum has been recorded after a thermal neutron irradiation with 4.3E16 cm^−2^ at the pneumatic rabbit facility RPA-3 of the FRM II research reactor and a counting time of 30 minutes one day after the end of exposure.

In conclusion, we could demonstrate that ionic signals exerted by NaCl had a strong impact on the polarization and enhancement of the Th2 program in human T helper cells. NaCl was able to replace IL-4 as the signal that was so far considered to have the strongest Th2 promoting effects for T cell polarization from naïve T cell precursors as well as for repolarization on the memory T cell level. NaCl accumulation in the human tissue might also promote the pathogenesis or increase the severity of allergic diseases as evidenced by highly increased sodium concentrations in the skin of patients suffering from atopic dermatitis. We therefore propose NaCl and its downstream signaling pathways as novel targets for the treatment of Th2-biased inflammatory diseases.

## Discussion

Adaptive immune cells integrate cues from the tissue microenvironment that tailor their phenotype and function for antigen-specific host defense and tolerance (*11, 32*). IL-4 has so far been considered a crucial cytokine for the final commitment and maintenance of Th2 immunity (*33*). We demonstrate here an IL-4-independent Th2 polarization pathway that is mediated by NaCl. NaCl acted both during the naïve T cell priming phase as well as at the memory T cell stage and could also divert alternative T helper cell fates into a Th2 phenotype. NaCl biased the Th1-Th2 dichotomy towards Th2 cell responses on multiple regulatory levels but also promoted effector Th17 cell responses. Our studies have identified NFAT-5 as well as its downstream target SGK-1 as molecular regulators of the Th2 cell program in hyperosmolar NaCl conditions. Importantly, the skin of patients with atopic dermatitis, a Th2 driven disease, was highly enriched in NaCl. Together, these results revealed NaCl as an ionic checkpoint for type 2 immunity and to be associated with the pathogenesis of atopic dermatitis. Therefore, targeting NaCl induced tonicity signaling could serve as a therapeutic strategy for the treatment of allergic diseases.

Previous studies demonstrated that NaCl promoted the differentiation of Th17 cells in polarizing cytokine conditions with implications for the pathogenesis of multiple sclerosis (*2–4*). Our data support and extend these conclusions not only on the naïve but also on the memory T cell level, even in the absence of exogenous polarizing cytokines. Although Th2-associated regulatory factors such as GATA-3 and Th2 effector cytokines were assessed in these previous studies as well, this occurred in Th17-polarizing cytokine conditions, which might have obscured the ability of NaCl to directly promote Th2 cells (*2*). In line with this reasoning, we did indeed observe abrogation of NaCl-induced IL-4 upregulation in human T cells under Th17 polarizing conditions. Furthermore, this finding suggests that the cytokine microenvironment modulates the impact of NaCl for T cell differentiation and function.

The dual support of NaCl for both Th2 and Th17 cell responses seems counterintuitive at first. The concomitant Th1/IFN-*γ* downregulation, however, supports the upregulation of IL-17 and IL-4 on the population level, which is known to be inhibited by the Th1 program (*34*). Our finding is in line with the previous observation that osmotic shrinkage of T cells by NaCl blunts IFN-*γ* expression (*35*). It does, however, challenge the recently propagated concept of salt-induced Th17 cell pathogenicity, which requires IL-17 to be coexpressed with IFN-*γ* on the single cell level (*2, 11, 36, 37*). Our data also demonstrate a strong degree of plasticity in effector T cells upon exposure to NaCl. IL-17 and IL-4 co-expression on the single-cell level could be induced in IL-17 producing T cells that were exposed to hypertonic NaCl concentrations. Of note, this hybrid IL-17 and IL-4 double-producing T cell population has previously been reported to be highly relevant in bronchial asthma and to arise from IL-17 single-positive T cells (*38*). Th1 cells were more resistant to IL-4 upregulation but downregulated IFN-*γ* in response to NaCl in support of the Th1 antagonistic effect of NaCl. Together, these results demonstrate that NaCl signaling skews the overall functionality of T cells towards the Th2 fate.

SGK-1 is a serine-threonine kinase that is activated by phosphorylation of its hydrophobic motif at Ser422 by the metabolic checkpoint kinase complex mTORC2 (*39*). mTORC2 therefore acts as an upstream activator of SGK-1. Our results are in accordance with previous reports that demonstrated a role for mTORC2 in the commitment to the Th2 lineage, but which missed to demonstrate which upstream signals differentially engaged the mTORC1 versus mTORC2 pathway (*40, 41*). It has been shown that T cell specific deletion of the mTORC2 specific adaptor Rictor abrogated the activation of SGK-1 (*40*). This resulted in reduced allergic asthma as well as increased anti-tumor and anti-viral immune responses in mouse models in accordance with the Th1-Th2 antagonism, which we demonstrated in T cells upon stimulation with NaCl. In addition, it has previously been shown that the Wnt antagonist Dickkopf-1 (DKK-1) promoted Th2 cell responses via the mTOR pathway and SGK-1 in mouse models of house dust mite-induced asthma or *Leishmania major* infection (*42*). Downstream targets of SGK-1 signaling following mTORC2 activation were shown to be JunB and the long isoform of the transcription factor TCF-1, for the Th2 versus Th1 fate, respectively (*39*). These previous insights into the control of T helper cell fates by mTOR complexes in mice corroborate our findings in the context of hypertonic NaCl induced T cell signaling. In particular, they propose NaCl as a novel, so far overlooked, upstream regulator of mTORC2 and thus of Th2 cell differentiation.

While the major transcriptional players that translate NaCl signaling into Th2 or Th17 cell responses have now been dissected, the upstream events of cellular sodium sensing still remain elusive. Recently, ion transport through the sodium-potassium-2 chloride cotransporter 1 (NKCC1) has been suggested to promote SGK-1 and IL-23R upregulation in Th17 cells in high-salt conditions (*43*). This effect was abrogated by specific pharmacological inhibitors of NKCC1, in particular by the diuretic drug furosemide, which supported the role of NKCC1 as the upstream sodium sensor on T cells. Together, these results imply that common diuretics may have modulatory effects on T cells in hyperosmotic tissue microenvironments and therefore possibly an impact on allergic diseases. In fact, furosemide has repeatedly been reported to improve allergic asthma by so far unknown mechanisms (*44, 45*). Its role in the NKCC1-SGK1-Th2 axis, however, remains to be investigated in detail to substantiate a therapeutic impact on the pathogenesis of Th2 mediated allergic diseases.

The conventional paradigm of salt homeostasis and systemic sodium equilibration by renal function has recently been refined. A new model has emerged by which NaCl concentrations are compartmentalized in distinct organs and tissues (*46*). Especially the skin has been reported to store sodium at concentrations exceeding those of blood and thus to display specialized, yet poorly defined mechanisms to store osmotically inactive NaCl (*47*). These insights warrant more investigations into the ionic composition of human organs and tissues. Our neutron activation analysis provides a tool that is highly sensitive and feasible for most human organs due to very small sample size requirements. It can therefore provide static information on the bodily distribution of many atomic elements of interest with high precision (*48*). Interestingly, we found highly elevated levels of sodium in the affected skin of atopic dermatitis patients (eczema) compared to unaffected control skin. This is in line with the Th2-mediated pathogenesis of this chronic inflammatory disease (*49*). Our finding that the cutaneous microenvironment of atopic dermatitis patients is highly enriched in sodium as well as the molecular dissection of NaCl-induced Th2 cell polarization could finally provide a mechanistic rationale for the early observational studies and therapeutic recommendations by Finkelstein. He published in his renowned pediatric reference book in 1912 that dietary salt restriction improves atopic dermatitis based on clinical observations (*7*).

Still the question remains of how and why sodium accumulates so drastically in atopic skin. Systemic abrogation of NaCl homeostasis can be excluded because serum concentrations of NaCl, which are controlled by renal function, are unaffected in atopic dermatitis. We consider a correlation of active inflammation with NaCl deposition unlikely due to lack of sodium deposition in other inflammatory skin diseases such as psoriasis. Atopic dermatitis is characterized by a profound barrier dysfunction. Continuous evaporation in the setting of this barrier defect promotes the local accumulation of NaCl in the skin (*50*). This is further supported by previous gene profiling studies, which reported the upregulation of genes encoding negatively charged glycosaminoglycans in lesional atopic skin (*51, 52*). These molecules are known to promote the accumulation of sodium ions in the skin by non-covalent binding (*53*). Interestingly, genes involved in glycosaminoglycan deposition (i.e. β1,3-Glucuronosyltransferase-I) are downstream targets of NFAT-5, which we have reported here to be the transcriptional regulator of the high-salt response (*54*).

An epithelial sodium sensor, Na_x_, has recently been shown to respond to perturbations of sodium homeostasis in the skin that is characterized by an atopic barrier defect (*55*). Although previous work has not investigated the direct link of Na_x_ with the immune system, sodium sensing has been implicated in fibroblast associated pathogenesis of AD. The *in vivo* knockdown of Nax in mice resulted in improvement of atopic dermatitis (*55*). Its downstream target ENaC, the major sodium channel in epithelial cells, is regulated by NFAT-5 as well as SGK-1, which we identified herein to regulate the Th2 bias (*50, 55, 56*). Together, these findings mechanistically link evidence of sodium deposition in the atopic skin with its impact on the pathogenesis of atopic dermatitis via the Th2-axis.

The profound barrier dysfunction in atopic dermatitis promotes the susceptibility to microbial invasion (*57, 58*). Previous studies have suggested that sodium might strengthen antimicrobial defense at barrier tissues (*59, 60*). Therefore, sodium accumulation in the atopic skin could represent a counter-regulatory microbial defense mechanism in the setting of genetic as well as Th2-mediated barrier dysfunction (*61*). In line with this reasoning, the atopic skin is indeed characterized by a microbial dysbalance (*62*). It is almost exclusively colonized by *S. aureus*, a bacterium which is known to be tolerant to high NaCl concentrations and to outgrow other bacteria in elevated NaCl conditions (*63, 64*). *S. aureus* skin colonization correlates significantly with the severity of atopic skin inflammation (*62*), whereas skin colonization with other bacteria is profoundly reduced in atopic dermatitis. Therefore, NaCl accumulation could provide a rationale for the microbial dysbiosis of atopic dermatitis, which has so far remained enigmatic. Taken together, a reductionist model could be envisioned for the pathogenesis of atopic dermatitis, which integrates the phenomenon of NaCl deposition in the atopic skin, the NaCl-induced Th2 bias of atopic immune deviation as well as the hegemony of NaCl resistant *S. aureus* (**Fig. S14**).

In conclusion, our investigations have revealed NaCl as an ionic checkpoint for human type 2 immunity with a potential clinical relevance for atopic dermatitis. Future studies into the compartmentalization and dynamic regulation of NaCl and other osmolytes in distinct human tissues will have to be conducted to unravel the full repertoire of tonicity-mediated immunomodulation in health and disease.

## Material and Methods

### Cell purification and sorting

Peripheral blood mononuclear cells (PBMC) were isolated by density gradient centrifugation using Ficoll-Paque Plus (GE Healthcare). CD4^+^ T cells were isolated from PBMC by positive selection with CD4-specific microbeads (Miltenyi Biotec) using an autoMACS Pro Separator. T helper cell subsets were sorted to at least 98% purity as follows: Th1 subset, CXCR3^+^CCR4^−^ CCR6^−^CD45RA^−^CD25^−^CD14^−^; Th2 subset, CXCR3^−^CCR4^−^CCR6^−^CD45RA^−^CD25^−^CD14^−^; Th17 subset, CXCR3^−^CCR4^−^CCR6^−^CD45RA^−^CD25^−^CD14^−^. Memory T helper cells were isolated as CD3^+^CD4^+^CD45RA^—^ lymphocytes, naïve T cells were isolated as CD45RA^+^CD45RO^−^ CCR7+ lymphocytes to a purity of over 98%. The antibodies used for sorting by flow cytometry were identical to those we have described previously (*11, 37*). Cells were sorted with a BD FACSAria III (BD Biosciences). Fresh peripheral blood was obtained from healthy donors. T cells from fresh healthy human skin (abdominoplasties) were isolated after 16 h digestion of the epidermis and dermis with collagenase IV (0.8 mg/ml, Gibco). Ethics approval was obtained from the Institutional Review Board of the Technical University of Munich (195/15s) and the Charité-Universitätsmedizin Berlin (EA1/221/11). All work was carried out in accordance with the Declaration of Helsinki for experiments involving humans.

### Cell culture

Human T cells were cultured in RPMI-1640 medium supplemented with 2 mM glutamine, 1 % (vol/vol) non-essential amino acids, 1 % (vol/vol) sodium pyruvate, penicillin (50 U/ml), streptomycin (50 μg/ml; all from Invitrogen) and 10 % (vol/vol) fetal calf serum (Biochrom). Hypersalinity (+NaCl) was induced by increasing NaCl concentrations by 50 mM NaCl compared to baseline cell culture medium including supplements (NaCl, Sigma-Aldrich). In some indicated experiments, T cell cultures were performed in the presence of recombinant cytokines (IL-6, 50 ng/ml; IL-12, 10 ng/ml; IL-4, 10 ng/ml; TGF-β, 5 ng/ml; IL-1β, 10 ng/ml; IL-23, 50 ng/ml; all from R&D Systems) or neutralizing antibodies (anti-IL-4, 10 μg/ml, BD Biosciences). Naïve T cells were labeled with carboxyfluorescein succinimidyl ester (CFSE) according to standard protocols and analyzed by gating on CFSE^low^ cells. T cells were stimulated with plate-bound anti-CD3 (2 μg/ml, clone TR66) and anti-CD28 (2 μg/ml CD28.2; BD Biosciences) for 48h before transfer into uncoated wells for another 3 days for a total culture period of 5 days unless indicated otherwise in the legends. T cell clones were generated in non-polarizing conditions as described previously following single-cell deposition with flow cytometry-assisted cell sorting or by limiting dilution (*65*). For experiments with murine T cells, single-cell suspensions were generated from spleen, inguinal, axial, and brachial lymph nodes of C57BL/6 and BALB/c mice. Mice were purchased from Janvier Labs and housed under specific pathogen-free conditions. All experiments were performed in accordance with the regulations of the Regierung von Oberbayern. Naïve CD4^+^ T cells were isolated by negative selection via the mouse naïve CD4^+^ T cell isolation kit (StemCell Technologies). Cells were seeded at a density of 200,000 cells/well in a flat-bottom 96-well plate, stimulated with plate-bound anti-CD3 and anti-CD28 antibodies (Th2 condition: 0.25μg/ml anti-CD3 and 2μg/ml anti-CD28; all other conditions 2 μg/ml anti-CD3 and 2ug/ml anti-CD28, BioXCell) and different mixtures of cytokines and monoclonal antibodies (Th0 condition: anti-IL-4, anti-IFN*γ*; Th1 condition: anti-IL-4, IL-12; Th2 condition: anti-IFN*γ*, IL-4; Treg condition: anti-IL-4, anti-IFN*γ*, TGF-β; Th17 conditions: anti-IL-4, anti-IFN*γ*, TGF-β, IL-6, or anti-IL-4, anti-IFN*γ*, TGF-β, IL-6, IL-23, or anti-IL-4, anti-IFN*γ*, IL-1β, IL-6, IL-23) with or without addition of 40 mM NaCl in DMEM (supplemented with 10% FCS, 10 mM HEPES, 1 mM sodium pyruvate, 1x MEM NEAA, 50 μM β-Mercaptoethanol, and 100 U PenStrep) (LifeTechnologies). Neutralizing antibodies and cytokines were used at the following concentrations: anti-IL-4 (10 μg/ml) and anti-IFN*γ* (10 μg/ml) from BioLegend; IL-4 (50 ng/ml), IL-12 (10 ng/ml), IL-6 (25 ng/ml), IL-1β (20 ng/ml), IL-23 (20 ng/ml) and TGF-β (2 ng/ml) from PeproTech. After 2.5 days in culture, cells were split into two new 96-well plates with a freshly prepared cytokine mix.

### Cytokine and transcription factor analyses

For intracellular cytokine staining, cells were restimulated for 5 h with PMA and ionomycin in the presence of brefeldin A for the final 2.5 h of culture (all Sigma-Aldrich). Cells were fixed and permeabilized with Cytofix/Cytoperm (BD Biosciences) according to the manufacturer’s instructions for intracellular cytokine analysis. For transcription factor analysis, cells were fixed and permeabilized with Cytofix/Cytoperm (eBioscience). Cells were stained with anticytokine antibodies (IL-4, IL-13, IFN-*γ*, IL-17A, IL-5; all from Biolegend) and with antibodies against transcription factors (GATA-3, ROR-*γ*t, both eBioscience; T-bet, Biolegend) or surface markers (IL-4R, R&D Systems; IL-12Rβ2, Miltenyi Biotec; CCR8, Biolegend) and were analyzed with a BD LSRFortessa (BD Biosciences), a CytoFLEX (Beckman Coulter) or a MACSQuant Analyzer (Miltenyi Biotec). Flow cytometry data were analyzed with FlowJo software (Tree Star) or with Cytobank (Cytobank Inc.). Cytokines in culture supernatants were measured by ELISA (R&D Systems) or by Luminex (eBioscience) according to standard protocols after restimulation of cultured T cells with phorbol-12-13-dibutyrate (50 nM, Sigma-Aldrich) and plate-bound anti-CD3 (1 μg/ml, TR66) for 8 h or as indicated in the respective figure legends. Counting beads (CountBright Absolute Counting beads, Thermo Fisher Scientific) were used to normalize for cell numbers if analysis of cumulative supernatants obtained from 5-day cell cultures was performed. For experiments with murine T cells, cells were analyzed by flow cytometry after a culture period of 3.5 days. Intracellular cytokine staining and staining for surface markers and transcription factors were performed as described previously (*66*). The following fluorochrome-conjugated monoclonal antibodies were used: anti-CD4 (clone RM4-5, Tonbo Biosciences); GATA-3 (TWAJ, eBioscience); IL-4 (11B11, BioLegend). Data were acquired on an LSRFortessa (BD Biosciences) or a BD FACSCanto II (BD Biosciences) and analyzed with FlowJo software (TreeStar).

For quantification of murine cytokine production by ELISA, cells were stimulated for 2.5 days *in vitro* in polarizing conditions, washed twice and resuspended in medium without any additional NaCl, cytokines, or blocking antibodies. After additional 24 h of incubation at 37°C, the supernatant was collected and the IL-4 concentration was determined by ELISA (Mouse IL-4 ELISA MAX™ Deluxe, BioLegend) according to the manufacturer’s instructions. Absorbance was measured with the GloMax^®^ Discover plate reader (Promega). IL-4 concentrations were normalized to the number of viable CD4^+^ T cells per well, which was determined by flow cytometry using counting beads (123count eBeads™ Counting Beads, Invitrogen).

### Gene expression analysis

The high capacity cDNA reverse transcription kit (Applied Biosystems) was used for cDNA synthesis according to the manufacturer’s protocol. Transcripts were quantified by real-time PCR with predesigned TaqMan Gene Expression Assays (SGK1, Hs00985033_g1; NFAT5, Hs00232437_m1; TBX21, Hs00203436_m1; GATA3, Hs00231122_m1; RORC2, Hs01076112_m1; 18S, Hs03928985) and reagents (Applied Biosystems). mRNA abundance was normalized to the amount of 18S rRNA and is expressed as arbitrary units (A.U.).

From 0.5 - 1 × 10^6^ snap-frozen cells total RNA was isolated for mRNA seq and DNaseI treated using the Direct-zol kit (Zymo Research) according to the manufacturer’s protocol. From ca. 200 ng total RNA, mRNA was bound to streptavidin coated tubes with the mRNA Capture kit (Roche). After three washes, first strand synthesis was performed with 200U M-MLV Reverse Transcriptase (Promega) at 37°C for 1.5 h. Second strands were generated using 10U DNA Polymerase I (Thermo Fisher Scientific), 200U T4-Ligase (NEB), 5U RNase H at 16° C for 2.5 h. Then cDNA was tagmented with 0.2 μl Tn5 enzyme from the Nextera Library Preparation kit (Illumina) for 5 min at 55° C. After clean-up with the MinElute PCR purification kit (Qiagen) and PCR amplification (9 - 12 cycles) with NEB Next PCR Master mix, samples were purified with AMPure XP beads (Beckman Coulter) and sequenced for 100 basepairs using a V3 single read flow cell on a HiSeq 2500 (Illumina). The generated data was trimmed for quality and adapter reads with TrimGalore! and then mapped with STAR aligner (*67, 68*). Duplicates were marked with the MarkDuplicates function from Picard tools (*69*). Reads were summarized with featureCounts. Differentially expressed genes (DEGs) were extracted using DeSeq2 package in R (*70*).

### Gene set enrichment analysis

Gene set enrichment was performed using GSEA v3.0 (http://www.gsea-msigdb.org/gsea/index.jsp). The differentially expressed genes from the transcriptome comparing CD4^+^CD45RA^−^ memory T cells cultured in low versus high NaCl conditions were ranked according to the Wald test statistics for differential expression provided by DESeq2 (RNA-Seq). The GSEA PreRanked tool was utilized in GSEA to compute the normalized enrichment score (NES). FDR q < 0.25 is considered a statistically significant enrichment. The NES relates to the enrichment score, which mirrors the degree to which a gene set is overrepresented at the top or bottom of a ranked list of genes. The Th1 and Th2 gene sets (gene set: GSE22886_Th1_vs_Th2 48h) were obtained from the Molecular signature database (MSigDB).

### Lentiviral gene silencing

Bacterial stocks containing lentiviral vectors with shRNA targeted against *SGK1* and *NFAT5* were purchased from Sigma-Aldrich (SGK1:TRCN0000196562, TRCN0000194957, TRCN0000040175, TRCN0000040177, TRCN0000010432; *NFAT5:* TRCN0000437810, TRCN0000020021, TRCN0000020022, TRCN0000020023, TRCN0000020019). All vectors were amplified and purified using the MaxiPrep or MidiPrep kit (Qiagen) according to the manufacturer’s instructions. Lentiviral particles were generated in HEK293 cells. 5 × 10^4^ presorted human memory T cells were activated on anti-CD3/CD28-coated plates for 12 h prior to transduction with supernatants from cultures with pooled lentiviral particles against *SGK1, NFAT5,* without shRNA insertion or with scrambled shRNA. After 48□h, cells were washed and selected with puromycin (Sigma-Aldrich). Gene expression was measured by quantitative real-time PCR 4 days after transduction.

### Targeted methylation analysis using NGS

Regional DNA methylation at genes of interest was analyzed using deep sequencing of bisulfite PCR amplicons. For PCR amplicon design, locus-specific primers for selected regions were designed using an in-house bisulfite primer design tool (oligos see **table S1**). At the 5‘ ends of each primer, NGS-compatible adapter tags were added (forward oligo tag: 5’TCTTTCCCTACACGACGCTCTTCCGATCT 3‘, reverse oligo tag: 5’GTGACTGGAGTTCAGACGTGTGCTCTTCCGATCT 3’). PCRs were set up in 30 μl reactions using 3 μl of 10x Hot Fire Pol Buffer (Solis BioDyne, Tartu, Estonia), 4 μl of 10 mM d’NTPs (Fisher Scientific, Pittsburgh, USA), 2.25 μl of 25 mM MgCl_2_ (Solis BioDyne, Tartu, Estonia), 0.6 μl of amplicon specific forward and reverse primer (10 μM each), 0.3 μl of Hot FirePol DNA Polymerase (5 U/μl; Solis Biodyne, Tartu, Estonia), 1 μl of bisulfite DNA and 18.25 μl of double distilled water. PCRs were run in an ABI Veriti thermo-cycler (Life Technologies, Karlsbad, USA) using the following program: 95°C for 10 min, then 40 cycles of 95°C for 1 min, 2.5 min of 56°C and 40 s at 72°C, followed by 7 min of 72°C and hold at 4°C. PCR products were cleaned up using Agencourt AMPure XP Beads (Beckman Coulter, Brea, USA). All amplified products were diluted to 4 nM. Next, amplicon NGS tags were finalized by PCR: a typical 50 μl reaction contained 25 μl of the DNA pool, 5 μl of 10x HotStartTaq buffer (Qiagen, Hilden, Germany), 4 μl of 10 mM d’NTPs, 2 μl of 25 mM MgCl_2_, 2.5 μl of 10 μM forward primer (5‘CAAGCAGAAGACGGCATACGAGATXXXXXXGTGACTGGAGTTCAGACGTGTGC TCTTCCGATCT 3‘, ,X’ marks a sample-specific barcode), 2.5 μl of 10 mM reverse-primer (5‘AATGATACGGCGACCACCGAGATCTACACXXXXXXTCTTTCCCTACACGACGCT CTTCCGATC 3‘, ,X‘ marks a sample-specific barcode), 0.6 μl of HotStartTaq polymerase (Qiagen, Hilden, Germany) and 8.6 μl of double distilled water. The reactions were incubated for 15 min at 97°C, followed by 5 cycles of 97°C (30 s), 60°C (30 s) and 72°C (30 s). After another ampure beads based clean up step, samples were quantified by a Qubit High Sensitivity Assay (Life Technologies, Karlsbad, USA) and diluted to 10 nM. Finally, all samples were pooled and loaded on an Illumina MiSeq sequencing machine. Amplicons were sequenced 2x250 bp (paired end) involving a MiSeq reagent kit V2 chemistry (Illumina, San Diego, USA). The raw sequencing data was quality checked using FastQC (v0.10.3; http://www.bioinformatics.babraham.ac.uk/projects/fastqc/) and trimmed for adaptors and low quality bases using the tools cutadapt (v1.3; https://code.google.com/p/cutadapt/) and Trim Galore! (v0.3.3; http://www.bioinformatics.babraham.ac.uk/projects/trim_galore/). Paired reads were joined using the FLASh tool (http://ccb.jhu.edu/software/FLASH/). Next, reads were sorted in a two-step procedure by (i) the NGS barcode adaptors to assign Sample ID and (ii) the initial 15 bp to assign amplicon ID. Subsequently, the sorted data was loaded into the BiQAnalyzer HT software 22 (http://biq-analyzer-ht.bioinf.mpi-inf.mpg.de/) using the following settings: the analyzed methylation context was set to “C”, minimal sequence identity was set to 0.9 and minimal conversion rate was set to 0.95. Only reads that covered more than 30% of all CpGs in an amplicon were used. The filtered high quality reads were then used for calling of methylation levels. Statistical analyses on obtained methylation calls were analyzed using in-house R-scripts.

### Quantification of skin sodium concentrations

Neutron activation analysis (NAA) at the Forschungsneutronenquelle Heinz Maier-Leibniz (FRM II) in Munich was employed as a non-destructive technique allowing the simultaneous detection of multiple chemical elements in 4 mm frozen punch biopsies from healthy human skin obtained during abdominoplasties or from atopic dermatitis patients (lesional and non-lesional). The emission of several gamma-quanta of characteristic energy upon decay of isotopes was measured following neutron-irradiation. The characteristic pattern of gamma radiation identified the emitting isotope. The intensity of the characteristic gamma radiation allowed the determination of the activity (given in Bq) of the emitter. The activity A at the end of the neutron exposure is connected to the number N of atoms of the mother isotope in the sample by means of the equation:

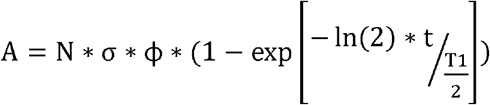

with □ being the thermal neutron flux density in 1/(cm^2^s), σ the activation cross section of the isotope in cm^2^, t the irradiation time and T_1/2_ the half-life of the radioisotope in equal units.

### Statistics

Student’s two-tailed paired t-test was used for statistical comparisons or as outlined in the respective figure legends; *p* values of 0.05 or less were considered significant. Analyses were performed using Prism 6 (GraphPad).

## Supplementary Material

**Figure S1. The viability of human T cells is preserved over a wide range of sodium chloride concentrations.** Human CD4^+^CD45RA^−^ memory T cells were sorted by flow cytometry and stimulated for 48h with CD3 and CD28 mAb for a total culture period of 5 days in the presence or absence of titrated concentrations of additional NaCl. T cell viability was assessed with Zombie fixable viability dyes and flow cytometry. Each circle represents an individual healthy blood donor and independent experiment.

**Figure S2. Th17 cell signature molecules are upregulated upon stimulation of memory T helper cells with sodium chloride.** (**A**) Human memory T helper cells were sorted from fresh PBMC as CD4^+^CD14^−^CD45RA^−^ T cells by flow cytometry and stimulated for a total culture period of 5 days in the presence (high) or absence (low) of additional 50 mM NaCl with CD3 and CD28 mAb for 48h. Intracellular staining and flow cytometry (FACS) on day 5 following PMA and ionomycin restimulation. Shown is one representative FACS staining (left panel) and cumulative data with each circle representing one donor. (**B**) T cells from three individual donors were isolated and stimulated as in (**A**) and then subjected to transcriptomic analysis by next-generation mRNA sequencing. Shown are boxplots of normalized gene expression levels for *IL26* (p=0.04). Red coloring denotes upregulation, blue coloring denotes downregulation.

**Figure S3. Sodium chloride induces the Th2 bias independently of exogenous or autocrine IL-4.** Human memory T helper cells were sorted from fresh PBMC as CD4^+^CD14^−^CD45RA^−^ T cells by flow cytometry and stimulated for a total culture period of 5 days in the presence (high) or absence (low) of additional 50 mM NaCl with CD3 and CD28 mAb for 48h. Anti-IL-4 neutralizing antibodies (10μg/ml) were added where indicated at the beginning of the culture. Intracellular staining and flow cytometry (FACS) on day 5 following PMA and ionomycin restimulation. Cytokine expression was normalized to control conditions (low NaCl). (n = 5-7)

**Figure S4. Sodium chloride induces upregulation of IL-4, IL-17 as well as IL-4 and IL-17 double positive T cells.** Human memory T helper cells were sorted from fresh PBMC as CD4^+^CD14^−^CD45RA^−^ T cells by flow cytometry and stimulated for a total culture period of 5 days in the presence (high) or absence (low) of additional 50 mM NaCl with CD3 and CD28 mAb for 48h. Intracellular staining and flow cytometry (FACS) on day 5 following PMA and ionomycin restimulation. Shown is one representative FACS staining of data shown in Fig.1A.

**Figure S5. Sodium chloride enhances Th2 and suppresses Th1 cell responses in T helper cell clones.** Human memory T helper cells were sorted from fresh PBMC as CD4^+^CD14^−^ CD45RA^−^ T cells by flow cytometry. T cell clones were generated from sorted cells during a 14 day culture period with irradiated allogeneic feeder cells and phytohemagglutinin before restimulation in low and high NaCl conditions with CD3 and CD28 mAb for 48h and a total restimulation period of 5 days. T cell clones were randomly selected from growing cultures for these restimulation experiments. Intracellular staining and flow cytometry (FACS) on day 5 following PMA and ionomycin restimulation. Shown is one representative FACS staining for each cytokine in a T cell clone (**A**) and cumulative data with each circle representing one individual T cell clone (**B**).

**Figure S6. Comparison of different osmolytes demonstrates sodium chloride to be the most potent inducer of IL-4 and suppressor of IFN-*γ* in memory T cells.** (**A**) Human CD4^+^ CD45RA^−^ memory T cells were sorted by flow cytometry and stimulated for 48h with CD3 and CD28 mAb for a total culture period of 5 days in the presence or absence of the indicated osmolytes (+ 50 mM NaCl, + 40 mM Na-Gluconate, + 30 mM MgCl2, + 80 mM mannitol, + 80 mM urea). Shown is one representative experiment (n, number of experiments, n = 3-6). (**B**) Human CD4^+^CD45RA^−^ memory T cells were stimulated as in (**A**) in the presence of a range of titrated concentrations of urea or mannitol, which did not adversely affect viability (viability ≥ 80%). (n=3)

**Figure S7. Memory CD8 T cells show stable IL-4 and IFN-*γ* expression upon stimulation with sodium chloride but upregulate IL-17.** CD8^+^CD45RA^−^ T cells were sorted by flow cytometry and stimulated for 48h with CD3 and CD28 mAb for a total culture period of 5 days in the presence or absence of 50 mM NaCl. Intracellular cytokine staining and flow cytometry on day 5 following restimulation for 5h with PMA and ionomycin.

**Figure S8. Skin T cells are distinct from blood T cells and differ in their expression of the tissue residency markers CD69 and CD103.** T cells were isolated by enzymatic digestion with collagenase IV from healthy human skin that was obtained from abdominoplasties. T cells were sorted by flow cytometry to obtain CD3CD56^−^ T cells and analyzed by flow cytometry for CD69 and CD103 expression. Shown is one representative experiment (n, number of experiments, n > 12).

**Figure S9. Sodium chloride induces the upregulation of the skin homing chemokine receptor CCR8, which enriches for Th2 associated cytokines.** Human memory T helper cells were sorted from fresh PBMC as CD4^+^CD14^−^CD45RA^−^ T cells by flow cytometry and stimulated for a total culture period of 5 days in the presence (high) or absence (low) of additional 50 mM NaCl with CD3 and CD28 mAb for 48h. (**A**) Flow cytometry (FACS) on day 5. Shown is one representative FACS staining (left panel) and cumulative data from individual experiments (right panel). n = 5. (**B**) CCR8^+^CD4^+^CD14^−^CD45RA^−^ and CCR8^−^ CD4^+^CD14^−^ CD45RA^−^ T cells (matched samples) were sorted by by flow cytometry and stimulated as in (**A**) including PMA and ionomycin restimulation for 5 hours. Intracellular cytokine staining and flow cytometry was performed. Each circle represents a separate blood donor.

**Figure S10. Th17 polarizing cytokine conditions abrogate the Th2 promoting effect of sodium chloride.** (**A**) Naïve CD4^+^CD45RA^+^CCR7^+^ T cells were sorted by flow cytometry and stimulated with the Th17 cell polarizing cytokines IL-1β, IL-6, TGF-β and IL-23 in the presence or absence of additional 50 mM NaCl for a total culture period of 5 days after stimulation with plate-bound CD3 and CD28 mAb for 48h. Intracellular cytokine staining and flow cytometry on day 5 following PMA and ionomycin restimulation. Shown is one representative experiment (n = 3). (**B**) Naïve CD4^+^CD45RA^+^CCR7^+^ T cells were stimulated as in (**A**) and supernatants were analyzed for IL-4 production by ELISA (n = 3, mean ± SEM).

**Figure S11. Mouse T cells differentiate into Th2 cells in response to sodium chloride.** Naïve murine CD4^+^ T cells were cultured *in vitro* under various T helper cell polarizing conditions with or without additional NaCl and analyzed on day 3.5 for IL-4 and GATA-3 expression by flow cytometry. (**A**) Representative histograms show intracellular IL-4 production after stimulation of cells with PMA and ionomycin. Cells treated with additional NaCl during the culture are depicted as black line and control cells are shown in grey. (B, C) Quantification of intracellular IL-4 production (as in A) and of GATA-3. *p < 0.05, **p < 0.01, ***p < 0.001, two-tailed paired t test (n = 5, mean ± SEM). (**D**) ELISA of cell culture supernatants harvested from Th2 polarizing T cell cultures (+IL-4) that were stimulated for 2.5 days in low or high NaCl conditions. Cells were replated in low NaCl conditions in the absence of exogenous cytokines after washing, and supernatants were harvested for analysis by ELISA after 24h. The cytokine concentration is normalized to 10.000 T cells for each condition. Each circle represents one mouse.

**Figure S12. SGK-1 is a downstream target of NFAT-5.** Human memory CD4^+^CD45RA^−^ T cells were stimulated for a total culture period of 5 days in the presence (high) or absence (low) of additional 50 mM NaCl with CD3 and CD28 mAb for 48h. SGK1 and NFAT5 expression were determined by αRT-PCR in the presence or absence of shRNA-mediated silencing of SGK1 and NFAT5 in low and high NaCl conditions (n = 4; low NaCl conditions, n = 9; high NaCl conditions; mean ± SEM). n.s., not significant (unpaired student’s t test).

**Figure S13. GATA-3 and T-bet are not regulated by NFAT-5 and SGK-1 in low sodium chloride conditions.** Human memory CD4^+^ CD45RA^−^ T cells were stimulated with CD3 and CD28 mAb for 48h and cultured for 5 days in the absence of additional NaCl (low NaCl). GATA3 and TBX21 expression were determined by αRT-PCR in the presence or absence of shRNAs targeting NFAT5 or SGK1.

**Figure S14. Model. Sodium chloride is highly enriched in the cutaneous microenvironment of atopic dermatitis skin lesions where it can promote Th2 cell responses.** Atopic dermatitis is characterized by an *S. aureus* dominated dysbiosis, which can also be promoted by NaCl, since *S. aureus* is tolerant to high NaCl concentrations in contrast to other species of the skin microbiota. Together, this model highlights a central role for NaCl as a novel microenvironmental factor of the atopic skin, because it links Th2 cell responses and dysbiosis as the main features of atopic dermatitis.

**Table S1. Selected regions and primer sequences used for DNA methylation analyses**

**Table S2. Patient information.** Clinical information on skin samples obtained from patients (**A**) with atopic dermatitis or psoriasis and from healthy control donors (**B**) including age, sex and disease activity (SCORAD, PASI). (**A**) and (**B**) show all parameters for each individual patient or donor. (**C**) Mean age for all included individuals. n.s., not significant.

## Acknowledgements

We thank Sophie Wolff for coordinating patient skin sample acquisitions and the irradiation at the neutron activation source in Garching. We thank Silke Hegenbarth for technical assistance and Christoph Heuser for critical review of the manuscript. We acknowledge the BMC Core Facility Flow Cytometry of the LMU Munich for providing equipment and the Flow Cytometry Core Facility of the German Rheumatism Research Center (DRFZ) in Berlin for cell sorting. Funding: This work was supported by the German Research Foundation (SFB1054 Teilprojekt B10 to C.E.Z and SFB1054 Teilprojekt B12 to D.B., Emmy Noether Programme BA 5132/1-1 to D.B.), the Fritz-Thyssen Stiftung (C.E.Z), and the German Center for Infection Research (C.E.Z.). Author contributions: JM (first author) performed most experiments, analyzed and interpreted the data. JM (co-author) performed all experiments with murine cells and together with DB analyzed and interpreted the data. RN and HM performed experiments and analyzed the data. HG, FJ, KE, DR and HM performed the measurements of sodium concentrations by neutron activation analysis, analyzed and interpreted the data. GG and JW performed the epigenetic experiments and analyzed and interpreted the data. GG and AW performed next-generation RNA sequencing. KN processed and analyzed the RNAseq data. SD and PK performed bioinformatic analyses with transcriptomic data sets. NGS, KE, TB and SG provided skin biopsies from atopic dermatitis patients and characterized and scored the patients samples. DS and YYC performed experiments with human T cells, analyzed and interpreted the data. CEZ conceived the study, supervised the experiments, interpreted the data and wrote the manuscript. All authors reviewed and approved the manuscript. Competing interests: The authors do not have any competing interests to declare.

## List of abbreviations

Th2: T helper 2 cells
IL: interleukin
PMA: phorbol myristate acetate
NaCl: Sodium chloride
n: number of experiments

